# Hippocampal network reorganization underlies the formation of a temporal association memory

**DOI:** 10.1101/613638

**Authors:** Mohsin S. Ahmed, James B. Priestley, Angel Castro, Fabio Stefanini, Elizabeth M. Balough, Erin Lavoie, Luca Mazzucato, Stefano Fusi, Attila Losonczy

## Abstract

Episodic memory requires linking events in time, a function dependent on the hippocampus. In “trace” fear conditioning, animals learn to associate a neutral cue with an aversive stimulus despite their separation in time by a delay period on the order of tens of seconds. But how this temporal association forms remains unclear. Here we use 2-photon calcium imaging to track neural population dynamics over the complete time-course of learning and show that, in contrast to previous theories, the hippocampus does not generate persistent activity to bridge the time delay. Instead, learning is concomitant with broad changes in the active neural population in CA1. While neural responses were highly stochastic in time, cue identity could be reliably read out from population activity rates over longer timescales after learning. These results question the ubiquity of neural sequences during temporal association learning, and suggest that trace fear conditioning relies on mechanisms that differ from persistent activity accounts of working memory.

## Introduction

Episodic memory recapitulates the sequential structure of events that unfold in space and time [Eichenbaum, 2017]. In the brain, the hippocampal network is critical for binding the representations of discontiguous events [Kitamura et al., 2015, Eichenbaum, 2017]. These findings are corroborated by recent evidence that the hippocampus generates sequences of neural activity that bridge the gap between sensory experiences [Pastalkova et al., 2008, MacDonald et al., 2011, Wang et al., 2015, Robinson et al., 2017], and that these dynamics are critical for memory [Wang et al., 2015, Robinson et al., 2017]. However, it remains a longstanding challenge to track how hippocampal coding is modified over the course of episodic learning that requires the association of events in time.

Pavlovian fear conditioning provides an attractive framework to study the neuronal correlates and mechanisms of associative learning in the brain [Letzkus et al., 2015, Grundemann and Luthi, 2015, Maddox et al., 2019, Grewe et al., 2017]. Classical “trace” fear conditioning (tFC) has long been used as a model behavior in the hippocampal literature for studying temporal association learning [Raybuck and Lattal, 2014, Kitamura et al., 2015]. In this paradigm, subjects learn that a neutral conditioned stimulus (CS) predicts an aversive, unconditioned stimulus (US), which follows the CS by a considerable time delay (the “trace” period). Circuitry within the dorsal hippocampus is required for forming these memories at trace intervals on the scale of tens of seconds [Raybuck and Lattal, 2014, Huerta et al., 2000, Quinn et al., 2005, Fendt et al., 2005, Chowdhury et al., 2005, Sellami et al., 2017]. Further, silencing activity in CA1, the output node of the hippocampus, during the trace period itself is sufficient to disrupt the temporal binding of the CS and US in memory [Sellami et al., 2017]. While these experiments pinpoint a role for hippocampal activity in forming trace fear memories, the underlying neural dynamics remain unresolved. Importantly, tFC precludes a simple Hebbian association of CS and US selective neural assemblies, due to the non-overlapping presentation of the stimuli.

Previous work has proposed that persistent activity mechanisms enable the hippocampus to connect representations of events in time, bridging time gaps on the order of tens of seconds. In particular, theories suggest that representation of the neutral CS and aversive US are linked through the generation of stereotyped, sequential activity in CA1 [Kitamura et al., 2015, Sellami et al., 2017], even when animals are immobilized [MacDonald et al., 2013]. Alternatively, hippocampal activity could generate a sustained response to the CS in order to maintain a static representation of the sensory cue in working memory over the trace interval, such as in attractor models of neocortical delay period activity [Amit and Brunel, 1997, Barak and Tsodyks, 2014, Takehara-Nishiuchi and McNaughton, 2008] and recently in the human hippocampus [Kaminski et al., 2017] during working memory tasks. However, these hypotheses of persistent activity remain to be tested during trace fear learning.

Here we leveraged 2-photon microscopy and functional calcium imaging to record the dynamics of CA1 neural populations longitudinally as animals underwent trace fear learning, in order to resolve the underlying patterns of network activity and their modifications in response to learning. Our findings show that persistent activity does not manifest during this paradigm, incongruous with sequence or attractor models of temporal association learning. Instead, learning instigated broad changes in network activity and the emergence of a sparse and temporally stochastic code for CS identities that was absent prior to conditioning. These findings suggest the role of the hippocampus in trace conditioning may be fundamentally different from learning that requires continual maintenance of sensory information in neuronal firing rates.

## Results

We previously developed a head-fixed variant of an auditory trace fear conditioning paradigm [Kaifosh et al., 2013], conducive to integration with 2-photon microscopy. Water-deprived mice were head-fixed and immobilized in a stationary chamber [Guo et al., 2014], to prevent locomotion from confounding learning strategy [MacDonald et al., 2013] (Fig. 1A). Mice were presented with a 20 sec auditory cue (CS), followed by a 15 sec temporal delay (“trace”), after which the animal received an aversive air-puff to the snout (US). A water port was accessible throughout each trial and we used animals’ lick suppression as a readout of learned fear [Kaifosh et al., 2013, Lovett-Barron et al., 2014, Rajasethupathy et al., 2015]. We first verified that learning in our head-fixed paradigm was dependent on activity in the dorsal hippocampus. Optogenetic inhibition of CA1 activity resulted in a significant reduction in lick suppression (Fig. S1), indicating that, as in freely moving conditions [Raybuck and Lattal, 2014, Huerta et al., 2000, Fendt et al., 2005, Chowdhury et al., 2005], head-fixed trace fear conditioning is dependent on the dorsal hippocampus.

**Figure 1.**
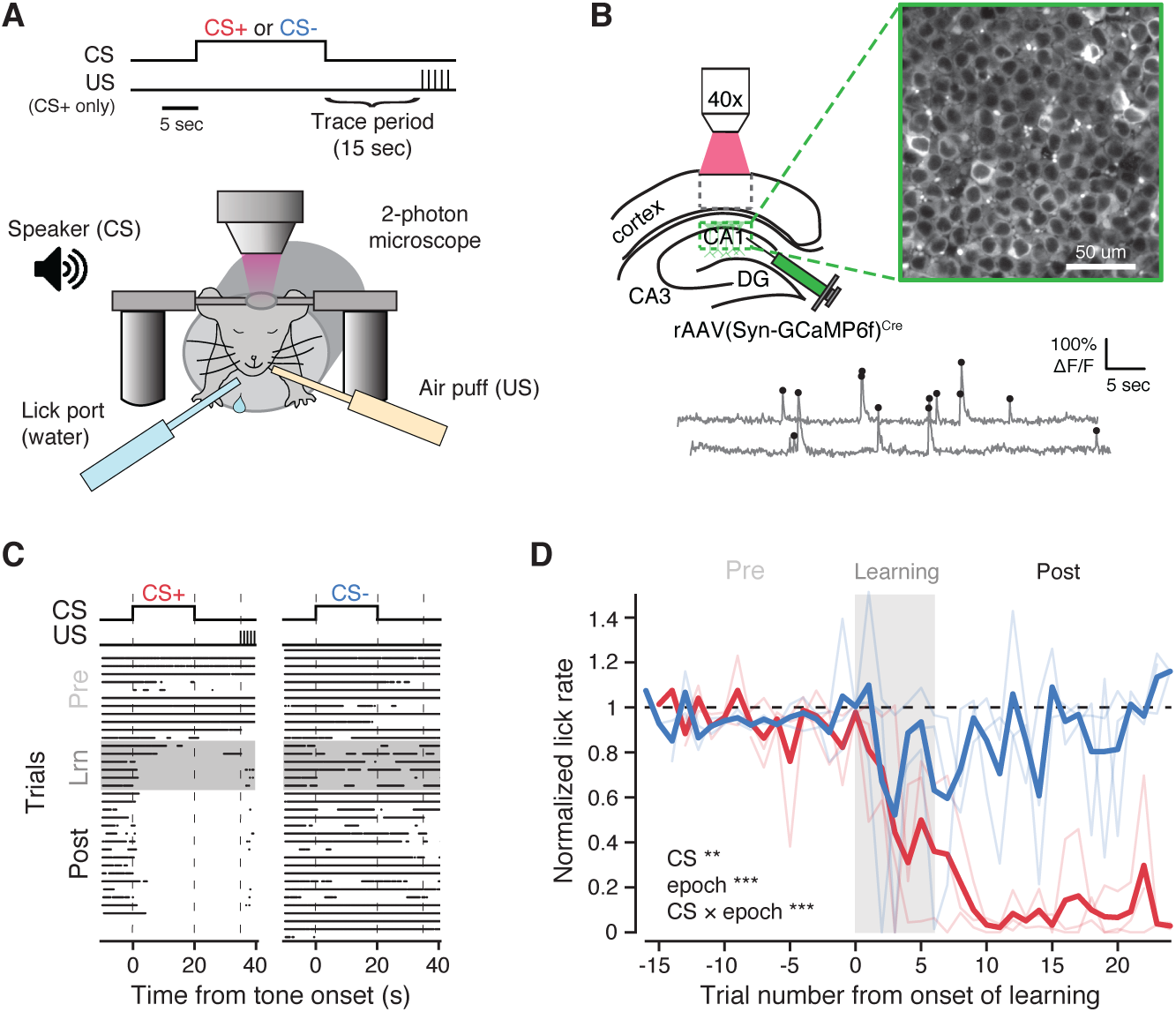
2-photon functional imaging of CA1 pyramidal neurons during differential trace fear conditioning. **A:** Schematic of the differential trace fear conditioning (tFC) paradigm. A head-fixed mouse is immobilized and on each trial exposed to an auditory cue (CS+ or CS-) for 20 seconds. This is followed by a 15-second stimulus-free ‘trace’ period, after which the US is triggered (CS+ trials only). Air-puffs are used as the US and lick suppression as a measure of learned fear. Operant water rewards are available throughout all trials. **B:** Top, schematic of *in vivo* imaging preparation with example 2-photon field of view in dorsal hippocampal area CA1. Bottom, calcium traces (grey) and inferred event times (black) from an example neuron. **C:** Behavioral data for an example mouse over the complete tFC paradigm. Each row is a trial, where dots indicate licks. CSs are first presented without US pairing (‘Pre-Learning’ epoch). Mice then rapidly learn to discriminate CSs and associate the CS+ with the US over the first 6 paired trials (‘Learning’ epoch), after which we continue to collect additional trials (‘Post-Learning’ epoch). **D:** Summary of behavioral dataset. We compute a normalized lick rate for each trial by dividing the lick rate during the CS tone (0-20 sec) period by the lick rate in the pre-CS (−10 to 0 sec) period. Bold lines are averages across mice. Thin lines show individual mice (n = 3 mice, linear mixed-effects model with fixed effects of CS and learning epoch, with mouse as random effect, main effect significance shown in inset, *post hoc* models fit to each epoch separately with fixed effects of CS and trial number, Pre-Learning: no significant effects, Learning: effect of trial number (***) and CS × trial number (*), Post-Learning, effect of CS (***), Wald χ^2^ test). \**p <* 0.05, \*\**p <* 0.01, \*\*\**p <* 0.001

In order to investigate the underlying network dynamics in the hippocampus that accompanied trace fear learning, we selectively expressed the fluorescent calcium indicator *GCaMP6f* in CA1 pyramidal neurons (Fig. 1B) via injection of a *Cre*-dependent AAV in *CaMKIIα-Cre* mice [Dragatsis and Zeitlin, 2000]. The head-fixation apparatus was mounted beneath the 2-photon microscope objective and mice were again water-restricted and trained to lick for water rewards while immobilized in the chamber [Guo et al., 2014]. Once mice licked reliably, we began neural recordings concurrent with a differential tFC paradigm (Fig. 1A), where on each trial mice were exposed to one of two auditory cues (CS+, CS-, either 10 or 2 kHz, identity randomized across mice). After collecting 10-15 “Pre-Learning” trials with exposure to each CS cue alone, we conditioned mice by selectively pairing CS+ trials with the US, delivered after the 15 sec trace interval. After the first 6 CS+/US pairings (“Learning” trials) wherein mice quickly acquired the trace association, we continued to record during 20-25 subsequent “Post-Learning” trials per cue with continued US reinforcement to avoid extinction of learned fear. Mice readily discriminated between the two cues throughout Post-Learning, as they suppressed licking consistently on CS+ trials but not CS-trials, where the air-puff was never presented (Fig. 1C,D).

Fluorescence imaging data from each trial was motion corrected [Kaifosh et al., 2014], and ROI spatial masks and activity traces were extracted using the Suite2p software package [Pachitariu et al., 2017]. All traces were deconvolved [Friedrich et al., 2017] to estimate underlying spike events. After registering ROIs across sessions, we identified 472 CA1 pyramidal neurons from 3 mice that were each active on at least 4 trials, which were used for subsequent analyses (Fig. 2A). Neural activity spanned all trial periods during the task both Pre- and Post-Learning, with a clear increase in neural activity following learning (population average event rate from 0-35 sec (*events/sec*): Pre: 0.039, Post: 0.057, *p <* 1.67e-10, signed rank test) and a large population response to the US (Fig. 2A).

**Figure 2.**
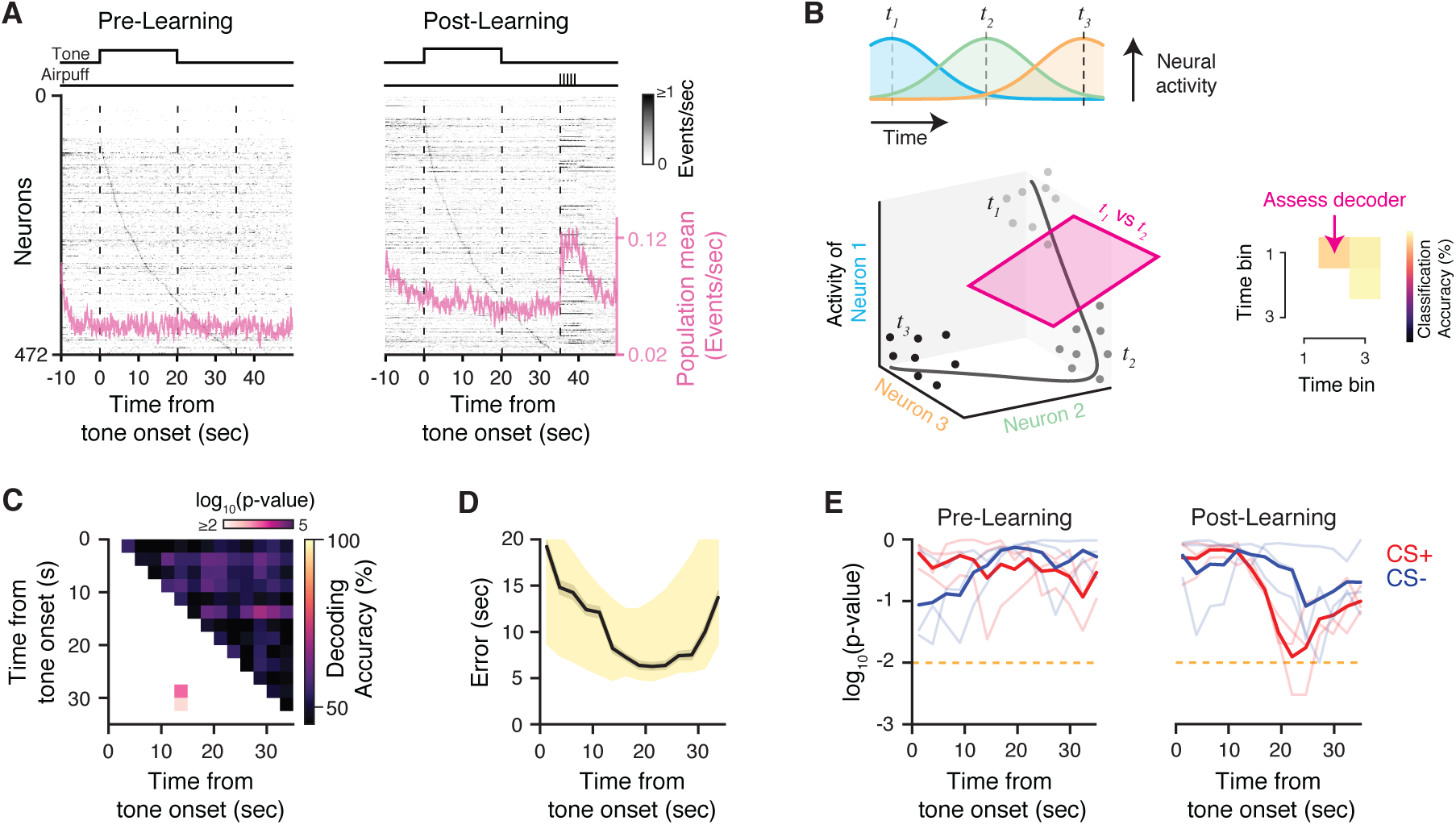
Temporal dynamics of CA1 population activity during trace fear conditioning. **A:** Summary of neural activity during Pre- and Post-Learning trials. For each epoch, activity is trial-averaged and neurons are sorted by the latency of their peak firing rate during the CS and trace periods (0-35 sec) during. The population average event rate is overlaid. **B:** Schematic of time decoding analysis. Top: trial-averaged tuning curves of a hypothetical sequence of time cells. Bottom: state space representation of the neural data. Dots indicate the neural state on single trials at three time points in the task. Right: A separate linear classifier (support vector machine; SVM) was trained to discriminate between population activity from every pair of time points in the task. **C:** Matrix of classifiers for an example mouse during Post-Learning CS+ trials. The upper triangle reports the cross-validated accuracy of classifiers trained to discriminate between the corresponding pair of time points. The lower triangle reports the p-value relative to a shuffle distribution. Most pairwise classifiers perform at chance level. **D:** Time prediction performance for the example shown in **C**. For each time bin in a test trial, the population activity is assessed by all classifiers, whose outputs are combined via a voting procedure to determine the decoded time. Decoding accuracy is assessed as the absolute error between real and predicted time. Black: cross-validated average and bootstrapped 95% confidence interval for time decoding error. Yellow shading: 95% bounds of the null distribution. Decoding error is within chance levels throughout the trial. **E:** Summary of time decoding significance relative to the null distribution during Pre-Learning (left) and Post-Learning (right) trials.

We first asked whether the hippocampus generated a consistent temporal code during each trial to connect the CS and US representations [Sellami et al., 2017, Kitamura et al., 2015]. While ordering population activity by the latency of neurons’ peak firing rates naturally lends the appearance of a sequence that spans the trial period (Fig. 2A), this ordering must be consistent across trials in order to be useful for computation. We approached this question through decoding, as the presence of sequential dynamics such as “time cells” [MacDonald et al., 2011] should allow us to decode the passage of time from the neural data [Bakhurin et al., 2017, Robinson et al., 2017, Cueva et al., 2019]. We used an ensemble of linear classifiers trained to discriminate the population activity between every pair of time points [Bakhurin et al., 2017, Cueva et al., 2019] in the tone and trace periods of the trial (0-35 sec, 2.5 sec bins). To illustrate the idea behind this analysis, we can summarize the activity of the network at each point in time as a point in a high dimensional neural state space, where the axes in this space corresponds to the activity rate of each neuron (schematized in Fig. 2B). The state of the network at each point in time during a trial traces out a path of points in neural state space. If the neural dynamics continually evolve in time (e.g. time cell sequences), then the neural state at one point in time (*t*) should be different from the states that occur at points further away in time (*t* + Δ*t*), reflecting the recruitment of different neurons at each point in the sequence. If these dynamics are reliable across many trials, we should be able to train a linear decoder to accurately classify whether data came from one time point or the other, by finding the hyperplane that maximally separates data from time *t* and *t* + Δ*t* in the neural state space. By extending this analysis to compare all possible pairs of time points (i.e. for all possible Δ*t*s), we can identify moments during the task that exhibit reliable temporal dynamics across trials [Bakhurin et al., 2017, Cueva et al., 2019]. We note however that the ability to decode time is not an exclusive feature of neural sequences, but a signature of any consistent dynamical trajectory where the neural states become sufficiently decorrelated in time (e.g. consider a population of “ramping” cells whose firing ranges change monotonically as a function of time [Cueva et al., 2019], or a chaotic trajectory in the activity space [Buonomano and Maass, 2009]). Accordingly, our analysis first addresses this broader question of whether *any* temporal coding arises during the task, without a priori assumptions on its parametric form.

We used these classifiers to assess whether neural activity was linearly separable between each pair of time points in the tone and trace periods of the task. Fig. 2C shows the results of this analysis for an example mouse during CS+ Post-Learning trials. The cross-validated decoding accuracy was no better than chance for most pairwise classifiers, suggesting that either the pattern of neural activity remains relatively constant throughout the tone and trace periods, or the dynamics are not consistent across trials. As an additional test of temporal coding, we can combine the output of the classifier ensemble to predict the time bin label of individual activity vectors from held out test trials (“one-vs-one” multi-class prediction, [Bakhurin et al., 2017, Cueva et al., 2019]). For each time bin in a test trial, the neural activity at that time is provided as input to all pairwise classifiers, whose binary decisions are combined via a voting procedure to determine the predicted time bin label of the data. Despite combining the information learned by all classifiers, time decoding accuracy did not exceed chance-level performance (Fig. 2D). We did not find evidence of significant temporal coding during either Pre- or Post-Learning trials (Fig. 2E); we did observe some trend toward significance during Post-Learning, which may reflect broader timescale differences in population activity during the CS and trace periods (see Fig. 4). Overall, these results suggest that CA1 neural activity sequences are not a reliable phenomenon during trace fear memory.

**Figure 3.**
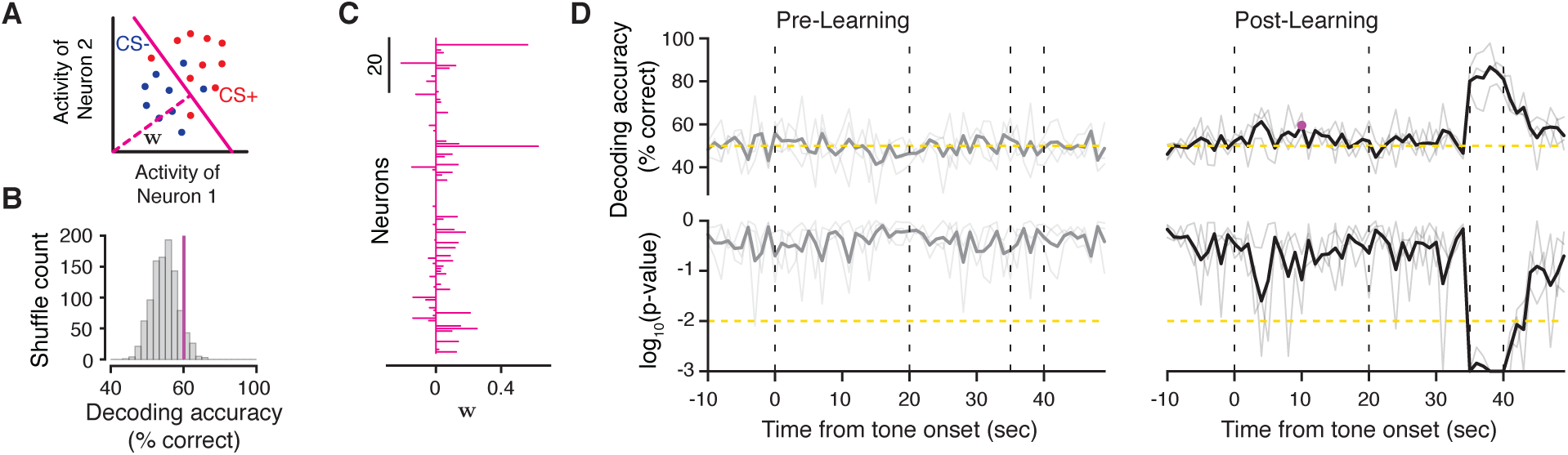
Stimulus identity is not persistently encoded in CA1. **A:** Schematic of CS decoding analysis. A separate classifier was trained to discriminate between CS+ vs CS-trials using population activity at each time point during the task (1 sec bins). **B:** Example CS decoder performance during Post-Learning. Purple line: average cross-validated decoding accuracy. Grey histogram: accuracy distribution obtained under surrogate datasets with shuffled trial labels. **C:** Population decoder weights for the classifier shown in **B** (averaged over cross-validation folds). **D:** CS decoding accuracy during Pre-Learning (left) and Post-Learning (right) trials, reported as % accuracy (top) and p-value relative to the null distributions (bottom). Instantaneous CS decoding accuracy is within chance levels throughout the trial, except during the time of air-puff delivery during Post-Learning trials. The example shown in **B** is marked in purple.

**Figure 4.**
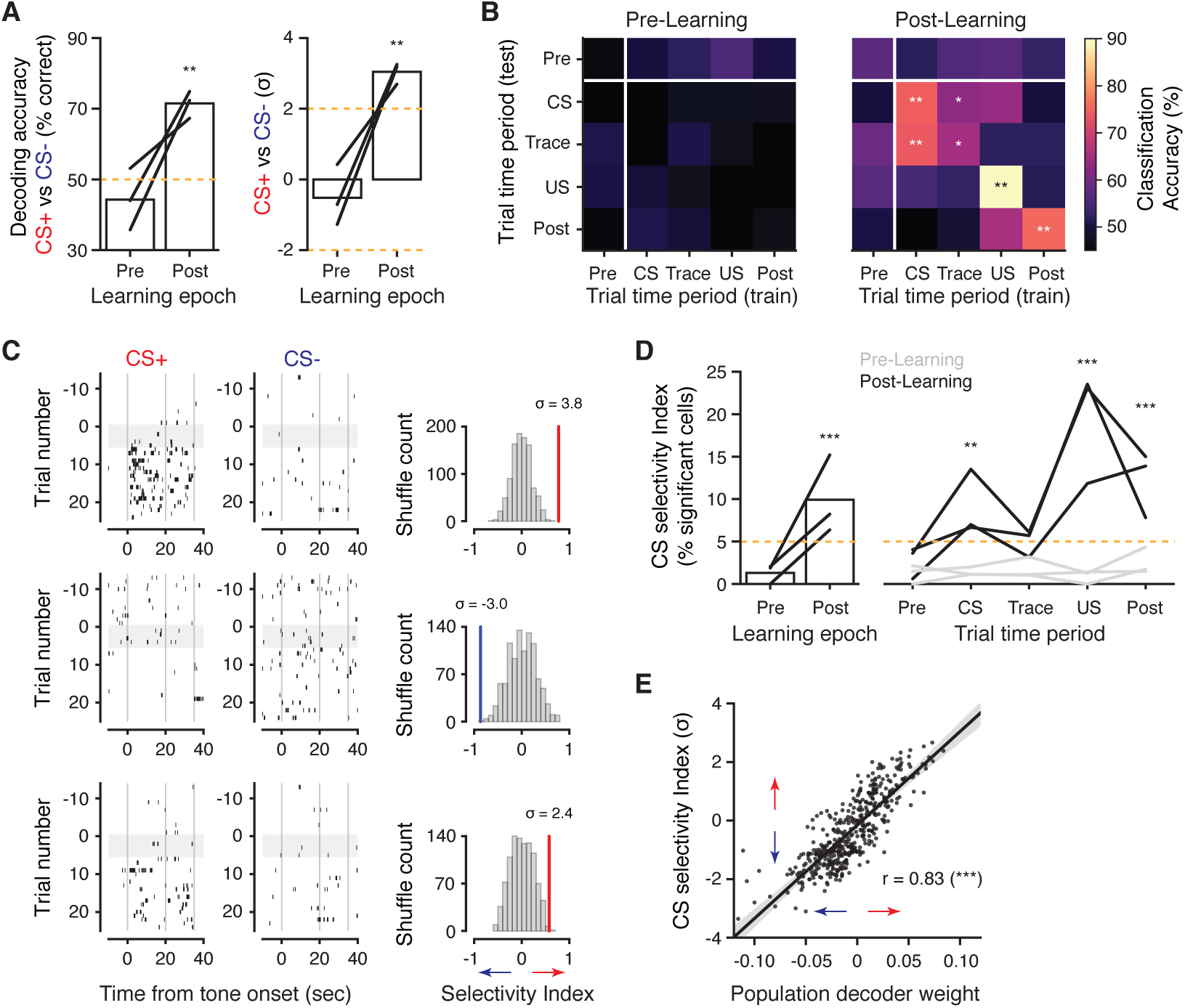
CS identity is predicted by CA1 activity rates on longer timescales. **A:** CS decoding accuracy for classifiers trained on the average activity within each trial’s CS and trace period. Left: % accuracy, right: z-normalized relative to null distributions calculated as in Fig. 3. Each line is the average cross-validated results from one mouse. **B:** Decoding CS identity from average activity in each trial time period. Decoders are trained and tested across each possible pair of time periods. In **A** and **B**, asterisks indicate significant p-values relative to shuffle distributions, averaged across mice. **C:** Raster plots of 3 simultaneously recorded CS-selective neurons (from average activity across CS and trace period). Right: Post-Learning CS selectivity index for each neuron, with shuffle distribution. **D:** Percentage of active cells with significant CS selectivity. Left: selectivity computed from average activity across CS and trace periods. Right: selectivity computed separately in each trial time period. Each line is a mouse. P-values indicate significant binomial test against the null hypothesis of *≤* 5%, pooled across mice. **E:** Regression of Post-Learning CS selectivity index for each neuron with its population decoder weight from **A.** Neurons with high selectivity are highly weighted by the population decoder (Pearson’s correlation). \**p <* 0.05, \*\**p <* 0.01, \*\*\**p <* 0.001

Since animals learned the association within the first few CS+/US pairings (Fig. 1C,D), we separately assessed whether any sequential dynamics might have rapidly and transiently emerged during the initial “Learning” trials. Due to the lower number of trials available, decoding was not possible, and so we computed a population sequence score by computing the rank correlation between the firing sequence of neurons across Learning trials (Fig. S2). However we found that neural activity did not organize into any reliable temporal patterns during these initial CS-US pairing trials. These data suggest that consistent temporal coding is not a dominant network phenomenon during trace fear conditioning, and so stereotyped sequential activity is unlikely to bridge the gap between CS and US presentations during the initial learning phase.

Our time decoding analyses indicated that most periods in time during the task were indistinguishable, which suggests that the network state during each trial may be relatively static. We considered an alternative hypothesis consistent with static activity, where CS information is maintained by persistent activation of a subgroup of hippocampal neurons [Kaminski et al., 2017], as in attractor models of neocortical networks suggested to underlie working memory [Amit and Brunel, 1997, Barak and Tsodyks, 2014, Takehara-Nishiuchi and McNaughton, 2008]. Under this scenario, the population dynamics would not evolve in time but discretely shift to a static state according to the trial’s CS cue, permitting the identity of the cue to be decoded throughout the duration of the trial. To test this, we trained a separate linear decoder at each time bin during the task to predict the identity of the CS cue. This analysis can be schematized as before, where the different CS cues are associated with different network states that can be reliably segregated in state space by a hyperplane (Fig. 3A).

For each time bin, we assessed whether we could accurately decode identity of the CS, separately analyzing Pre- and Post-Learning trials. Fig. 3B show the results of this analysis for an example Post-Learning time bin, reported as the percentage of correctly decoded trials and compared to a null distribution generated by shuffling the trial type identities. Our choice of linear classifiers also allowed us to obtain an intuitive measure of the importance of each neuron to the decoder’s decisions, by examining the weights of each neuron along a vector orthogonal to the separating hyperplane (**w**, Fig. 3A,C, Stefanini et al. [2019]). Examining these results across time bins, we found that we were unable to decode the CS identity during Pre-Learning or Post-Learning trials at any point prior to the delivery of the US (Fig. 3D). We verified that this was not due to our choice of decoder; CS decoding conclusions were unchanged when we used a Bayesian approach (Fig. S3). Similar methods have been used previously during appetitive trace conditioning paradigms in monkeys, where the CS identity and US prediction could be accurately decoded from the moment-to-moment population activity in the amygdala and prefrontal cortex [Saez et al., 2015]. However, in these experiments the trace period was an order of magnitude shorter than in our fear learning paradigm. These results indicate that information about the CS identity does not appear to be maintained in the moment-to-moment activity of CA1 pyramidal cell populations. Relatedly, activity was not robustly tied to instantaneous licking behavior, which differed markedly between cues during Post-Learning trials (Fig. 1D). CS decoding accuracy was generally high during US-delivery in Post-Learning trials, consistent with the clear population response to the air-puff (Fig. 2A). Though still within chance-level, there was a visible increase in the variability of classifiers’ performance during the tone delivery in Post-Learning trials. This suggested to us that there may be cue-selective responses in the population that appeared with variable timing across trials, and so they could not be reliably decoded at more granular time resolutions.

We tested the hypothesis that neural activity levels were predictive at longer timescales, first by attempting to decode the CS identity from the average activity rate across the CS and trace periods of each trial (Fig. 4A). This analysis is identical to that outlined in Fig. 3, except we collapsed the activity into a single time bin by averaging each neuron’s activity trace from 0-35 sec. Surprisingly, we found that at this timescale, we could significantly decode the CS identity from the population activity rates. Decoding accuracy exceeded chance performance only during Post-Learning trials, congruent with a change in network organization following learning.

As external sensory information about the stimulus is only available during the 20 sec CS period, we next asked whether average activity rates during other trial periods were still predictive of the cue identity, and how those activity patterns compared to those present during the tone presentation. To address this, we constructed a cross-time period decoding matrix, where we trained decoders to predict the CS identity using the average activity in a given time block and tested their performance on the activity from all trial time blocks (Fig. 4B). Values along the diagonal of the matrix gauge how reliably activity during each time block predicts the cue, while off-diagonal entries assess how decoders generalize to other trial periods. As expected from Fig. 4A, decoding during Pre-Learning trials was at chance level for all conditions. During Post-Learning, significant decoding accuracy was observed in all time blocks starting from the tone onset (Fig. 4B). Additionally, decoders showed significant generalization between the CS and trace periods, suggesting that a representation of the tone is maintained in the stimulus-free trace interval, and that this representation is highly similar to activity during stimulus presentation itself. Decoders trained on the trace period performed notably worse than those trained on the CS period, whether tested on the trace or CS period, indicating that activity during the stimulus-free trace period was less stable than in the CS period. This analysis also showed that the representation of the CS and US were largely distinct following learning, unlike observations in the basolateral amygdala during associative learning [Grewe et al., 2017].

Our analysis established that stimulus identity could be read out from the population activity during the tone and trace period in a learning-dependent manner, and so we sought to connect these findings to changes in neural activity at the level of individual neurons. While some neurons exhibited very robust cue preferences following learning (Fig. 4C, top), these were rare and most cells showed more graded firing rate changes (Fig. 4C, bottom). We quantified single neuron tuning via a selectivity index, standardized against a shuffle distribution generated by shuffling trial type identities, and measured the fraction of significantly CS-tuned neurons. Single neuron tuning heavily mirrored the population decoding results, both when computed over the tone and trace periods combined and in individual trial time periods (Fig. 4D). Similar to the drop in decoding accuracy, the fraction of tuned neurons was lower during the trace period than the CS period for all mice. Overall, neurons’ normalized CS selectivity indices were extremely correlated with their weight in the population decoder (Fig. 4E), demonstrating that the decoding analysis most heavily relied on neurons in the population with strong tuning to CS identity.

Finally, we sought to characterize how network structure changed during task learning on a trial-to-trial basis. Across the different learning epochs of the task, the fraction of active neurons in the population significantly increased (Fig. 5A). In addition to heightened network activity, we asked how the set of active neurons compared across trials. To address this question, we measured the overlap between the set of neurons active during the CS and trace periods between each pair of trials using the Jaccard similarity index (Fig. 5B). These scores were standardized by a null distribution generated by shuffling the active neuron pool in each trial, in order to control for differences driven trivially by disparities in the number of active neurons. Fig. 5C depicts the trial-by-trial overlap in neural ensembles for an example mouse for the complete time course of the experiment, revealing a marked shift in activity patterns from Pre- to Post-Learning. To summarize these observations, we computed the average ensemble similarity to Pre- or Post-Learning trials for each trial in the experiment (Fig. 5D). These results demonstrate that trace fear learning is accompanied by a large modification in the active neuronal population, beyond that expected from the overall increase in network activity following conditioning. Given the established role of hippocampal circuits in contextual memory formation [Maren et al., 2013, Urcelay and Miller, 2014, Fanselow, 2010], this change may also reflect a learning-dependent change in the representation of the broader context, in addition to or independent of the encoding of the individual CS cues.

**Figure 5.**
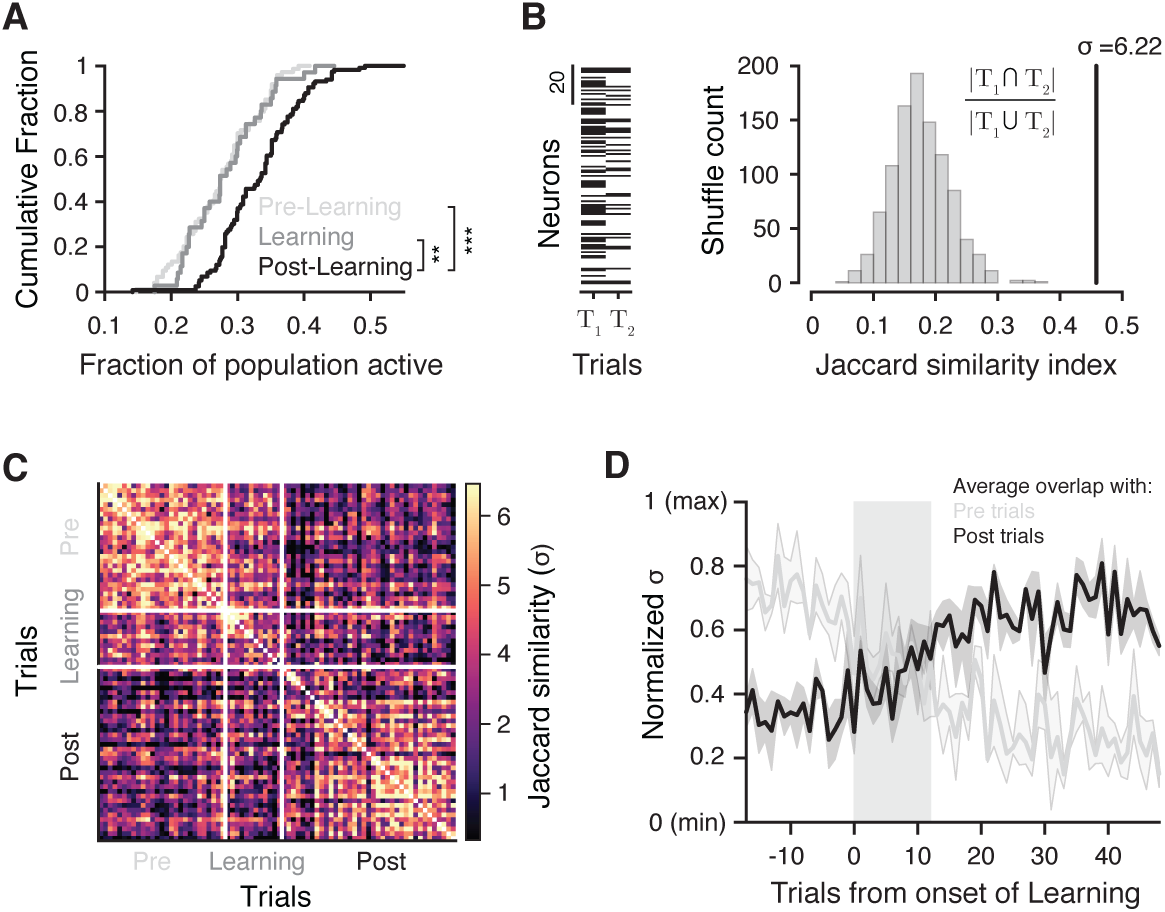
Learning parallels a shift in the active neural population. **A.** Fraction of neurons active during the tone and trace period, per trial (Kolmogorov-Smirnov test). Trials for both CS cues are pooled. **B.** Left: example binary activity vectors for two trials. Black indicates the neuron was active on that trial. Right: Jaccard similiarity index for the trial pair shown at left. The observed similarity index is z-scored to a null distribution generated by shuffling the neuron identities on each trial, to control for differences in the number of active neurons. **C:** Matrix of pairwise z-scored trial similarities for an example mouse. **D:** Average similarity to Pre-(grey) and Post-Learning (black) trials, for each trial in the experiment (excluding trial self-comparisons). Each mouse is first normalized by its minimum and maximum similarity scores. Learning trials are shaded in Grey. The set of active neurons changes during Learning. \**p <* 0.05, \*\**p <* 0.01, \*\*\**p <* 0.001

## Discussion

Here we have implemented a novel experimental framework for deciphering neural coding during non-spatial, temporal associative learning in the hippocampus using chronic cellular imaging. These methods demonstrate that network dynamics during trace fear conditioning are inconsistent with hypotheses of persistent sequential [Kitamura et al., 2015, Sellami et al., 2017] or sustained [Kaminski et al., 2017] activity in CA1. Rather we find that behavioral learning in CA1 is underpinned by the emergence of a subset of cue-selective neurons with stochastic temporal dynamics across trials. These units encode cue information in a learning-dependent manner. It is plausible that these dynamics may relate to descriptions of hippocampal memory “engram cells”, identified via immediate early gene (IEG) products [Liu et al., 2012, Vetere et al., 2017, Tanaka et al., 2018, Rao-Ruiz et al., 2019].

Consistent with this notion, our estimates of significantly CS-selective cells that emerge with learning fall within the range of the 10-20% of CA1 pyramidal neurons recruited in engrams supporting hippocampal-dependent memory [Tayler et al., 2013, Tanaka et al., 2018, Rao-Ruiz et al., 2019]. If the sparse subset of CS-selective cells we observe do represent engram cells, then this would further support the notion that gating mechanisms exist to ensure sparsity of encoded engrams, since the size of the engram does not vary with behavioral paradigms, such as immobilized, auditory tFC used here compared with contextual learning in freely moving animals [Rao-Ruiz et al., 2019]. It is important to note however that in our data, CS-selectivity at the single cell level manifested along a continuum of firing rate differences between conditions (Fig. 4), and it is unclear how coding differences at these scales would be resolved with IEG-based methods.

Our data show that, prior to and following learning, cue information is not actively transmitted by neurons’ moment-to-moment firing rates. Neural activity is instead remarkably sparse across time and conditions. The lack of consistent CS coding during the Pre-Learning epoch is consistent with prior evidence that few CA1 pyramidal neurons respond to passive playback of auditory stimuli [Aronov et al., 2017]. It is possible that these dynamics also differ according to sensory modality and behavioral states, such as locomotion. In previous studies that report neural sequences in CA1 during delay periods [Pastalkova et al., 2008, MacDonald et al., 2011, Wang et al., 2015, Robinson et al., 2017], the hippocampal network state was in a regime largely dominated by strong theta oscillations in local field potentials (LFP) and frequent burst firing by pyramidal neurons [Buzsáki and Moser, 2013] reminiscent of activity during active behaviors such as spatial exploration. The dynamics we observe resemble more closely the activity often seen during immobility and awake quiescence, where pyramidal neurons fire only sparsely and in a manner often restricted to population bursts associated with sharp-wave ripple (SWR) LFP events [Buzsáki, 2015].

We observe sparse and temporally variable activity that nevertheless is predictive of task information when averaged over longer time periods. It is possible then that these dynamics may arise from stochastic reactivation of memory traces via SWR events. Given the general sparsity of activity during the task and the unreliability of single spike detection with calcium imaging, it is likely that we have underestimated the task-related activity here. Detection of population burst events is possible at faster imaging speeds and with dense sampling of CA1 populations [Malvache et al., 2016], and we speculate that these events may be the underlying cause of the detected CS information following learning. This idea could connect the separate observations that inactivation of the medial entorhinal cortex, a major input to the hippocampus, disrupts both the structure of SWR events in CA1 [Yamamoto and Tonegawa, 2017] and trace fear conditioning [Suh et al., 2011].

Sparse reactivation of neural assemblies may also suggest a fundamentally different mode of propagating information over time delays during trace fear learning, for example, by storing information transiently in synaptic weights [Mongillo et al., 2008, Barak and Tsodyks, 2014]. Such a method could confer a considerable metabolic advantage for maintaining memory traces over long time delays, in contrast to generating persistent activity or neural sequences. Previous theoretical work to this end has focused on short-term plasticity in networks with pre-existing attractor architectures. There, pre-synaptic facilitation among the neurons in a selected attractor drives its reactivation in response to spontaneous input [Mongillo et al., 2008], by out-competing the other attractors. To preserve information about the reactivated attractor on long timescales, they propose a mechanism involving some form of ongoing refreshing activity to maintain synaptic facilitation, which would be incompatible with our observations. As a consequence, the time constant of facilitation limits the lifetime of these memory traces to around the order of a second, which is much shorter than the trace period we considered here. Alternatively, we speculate that coding assemblies may develop through continual Hebbian synaptic potentiation over trials, and that plasticity induced by the most recently presented cue may provide a bias in reactivated network states on each trial by increasing the depth of the corresponding basins of attraction. A similar scheme has been explicitly modeled in the case of visuo-motor associations [Fusi et al., 2007] and it is known to require synaptic modifications on multiple timescales [Benna and Fusi, 2016]. However, it has never been considered in the case of fear conditioning and long time intervals and will be an important direction for future work.

Though differential neural responses to the cues tended to be subtle, we observed an overall marked turnover in the set of active neurons from Pre- to Post-Learning trials that was common for both CS+ and CS-trials. It is possible that this is a broader change in the hippocampal representation that associates the context with the US itself, or reflects an association with more abstract knowledge of the cue-outcome rules [Maren et al., 2013, Urcelay and Miller, 2014, Fanselow, 2010]. Relatedly, past work has shown that memories experienced closely in time may be encoded by overlapping populations of neurons [Cai et al., 2016], consistent with our findings of a largely shared neural ensemble between CS trial types in Post-Learning. This linking of distinct but related memories to overlapping populations may occur as a result of transient increases in the excitability of neural subpopulations, which biases allocation into memory engrams [Yiu et al., 2014, Cai et al., 2016, Rashid et al., 2016].

Our findings highlight a hippocampal-dependent learning process that associates events separated in time in the absence of persistent activity. Given that associations in real-world scenarios are often dissociated from emotionally valent outcomes by appreciable time delays [Raybuck and Lattal, 2014], our findings have broad implications for models of temporal association learning and circuit dynamics underlying the dysregulation of anxiety and fear in neuropsychiatric disorders.

## Acknowledgments

We thank S.A. Siegelbaum, R. Hen, P.D. Balsam, and C.D. Salzman for fruitful discussions. We thank B.V. Zemelman for viral reagents used in this study. This work was supported by Leon Levy Neuroscience and American Psychiatric Foundation / Genentech, Inc Research Fellowship Awards (to MSA), NIH grants (R25MH086466, T32MH018870, and K08MH113036 to MSA; T32NS064928 to JBP; K25DC013557 to LM; R01MH100631, U19NS090583, and R01NS094668 to AL), NSF’s NeuroNex program award DBI-1707398 (to FS and SF), the Simons, Grossman, and Gatsby Charitable Foundations (to FS and SF), the Kavli Foundation (to FS, SF, and AL), the Searle Scholars and Human Frontier Science Programs (to AL), and the McKnight Memory and Cognitive Disorders Award (to AL).

## Author Contributions

M.S.A., J.B.P., S.F., and A.L. conceived the project. M.S.A. conducted experiments with inputs from J.B.P., L.M., S.F., and A.L. J.B.P. developed and performed data analyses with L.M., F.S., and S.F., with inputs from M.S.A. and A.L. A.C. designed behavioral hardware. A.C., E.M.B., and E.L. provided technical support. M.S.A, J.B.P., L.M., S.F., and A.L. wrote the manuscript, with input from all authors.

## Materials and Methods

### Behavior and Imaging

#### Mice and viruses

All experiments were conducted in accordance with the NIH guidelines and with the approval of the Columbia University Institutional Animal Care and Use Committee. Optogenetic experiments were performed with adult (8-14 weeks) male and female C57/Bl6 mice (Jackson Laboratory) injected bilaterally (see below) with either recombinant adeno-associated virus (rAAV) expressing *ArchT* (rAAV1/2-*Syn-ArchT*) or *tdTomato* control protein (rAAV1/2-*Syn-tdTom*), under the Synapsin promoter. These viruses were the generous gift of Dr. Boris Zemelman. Imaging experiments were performed with adult (8-16 weeks) male and female transgenic *CaMKIIα-Cre* mice on a C57/Bl6 background, where *Cre* is predominantly expressed in pyramidal neurons (R4Ag11 line, Dragatsis and Zeitlin [2000]; Jackson Laboratory, Stock No: 027400). *Cre*-dependent recombinant adeno-associated virus (rAAV) expressing *GCaMP6f* (rAAV1-*Syn-Flex-GCaMP6f-WPRE-SV40*, Penn Vector Core) was used for labeling of pyramidal neurons.

#### Surgical procedure

Viral delivery to hippocampal area CA1 and implantation of headposts, optical fibers, and imaging cannulae were as described previously [Kaifosh et al., 2013, Kheirbek et al., 2013, Lovett-Barron et al., 2014]. Briefly, viruses were delivered to dorsal CA1 by stereotactically injecting 50 nL (10 nL pulses) of rAAVs at three dorsoventral locations using a Nanoject syringe (−2.3 mm AP; −1.5 mm ML; −0.9, −1.05 and −1.2 mm DV relative to bregma). For head-fixed optogenetic experiments, mice were chronically implanted with bilateral optical fiber cannulae above the CA1 injection sites after virus delivery [Lovett-Barron et al., 2014, Kheirbek et al., 2013]. A stainless steel headpost was then fixed to the skull [Kaifosh et al., 2013]. The cannula, headpost, and any exposed skull were secured and covered with black grip cement to block light from the implanted optical fibers. For imaging experiments, mice were allowed to recover in their home cage for 3 days following virus delivery procedures. They were then surgically implanted with a steel headpost along with an imaging window (diameter, 3.0 mm; height, 1.5 mm) over the left dorsal hippocampus. Imaging cannulae were constructed by adhering (Narland optical adhesive) a 3 mm glass coverslip (64-0720, Warner) to a cylindrical steel cannula. The imaging window surgical procedure was performed as detailed previously [Kaifosh et al., 2013, Lovett-Barron et al., 2014]. For all surgeries, analgesia was continued for 3 days postoperatively.

#### Behavioral apparatus

We adopted our previously described [Kaifosh et al., 2013, Lovett-Barron et al., 2014] head-fixed system for combining 2-photon imaging with microcontroller-driven (Arduino) stimulus presentation and behavioral read-out. To maintain immobility and constrain neural activity related to locomotion [MacDonald et al., 2013], mice were head-fixed in a body tube chamber [Guo et al., 2014]. The chamber was lined with textured fabric that was interchanged between trials to prevent contextual conditioning. Tones were presented via nearby speakers and air-puffs delivered by actuating a solenoid valve, which gated airflow from a compressed air tank to a pipette tip pointed at the mouse’s snout. Water reward delivery during licking behavior was gated by another solenoid valve in response to tongue contact with a metal water port coupled to a capacitive sensor. Electrical signals encoding mouse behavior and stimulus presentation were collected with an analog-to-digital converter, which was synchronized with either optogenetic laser delivery or 2-photon image acquisition by a common trigger pulse.

#### Head-fixed trace fear conditioning

Starting 3-7 days after surgical implantation, mice were habituated to handling and head-fixation as previously described [Kaifosh et al., 2013, Lovett-Barron et al., 2014, Guo et al., 2014]. Within 3 days, mice could undergo up to an hour of head-fixation on the behavioral apparatus while remaining calm and alert. They were then water deprived to 85-90% of their starting body weight and trained to lick operantly for small-volume water rewards (500 nL/lick) while head-fixed. Before undergoing experimental paradigms, mice were required to maintain consistent licking for multiple (6-12) 60 second trials per day while maintaining their body weight between 85-90% of starting weight.

For optogenetic experiments, we utilized our previously described head-fixed ‘trace’ fear conditioning paradigm [Kaifosh et al., 2013]. Briefly, we paired a 20 second auditory conditioned stimulus (CS, either 10 kHz constant tone or 2 kHz tone pulsed at 1Hz) with air-puffs (unconditioned stimulus, US; 200 ms, 5 puffs at 1 Hz), separated by a 15 second stimulus-free ‘trace’ period. During each conditioning trial, we recorded licking from mice over a 55 second period: 10 second pre-CS, 20 second CS, 15 second trace, and 5 second US. Mice were conditioned across trials spaced throughout three consecutive days. On each trial, we used suppression of licking during the tone, normalized to licking during the 10 second pre-CS period, as a measure of conditioned fear. We changed the fabric material in the behavioral chamber between every trial to prevent contextual fear conditioning [Kaifosh et al., 2013, Lovett-Barron et al., 2014].

For 2-photon imaging experiments, we expanded our behavioral paradigm to a differential learning assay using the 2 different auditory cues above as either a CS+ or CS-(where only CS+ is paired with the aversive US). We randomized the assignment of CS+ and CS-tones across mice. Prior to the introduction of US-paired conditioning trials, we obtained multiple trials of behavioral responses (10-15 trials; “Pre-Learning”) to each CS cue presented alone in pseudorandom order over 3 days. This helped to confirm that subsequent lick suppression was not due to ‘pseudoconditioning’ or stimulus novelty effects [Burman et al., 2014]. We then subjected mice to our 3-day conditioning protocol with US-pairing as above, but with alternation between CS+ and CS-trials (‘Learning’). Finally, over another 3 days, we collected additional trials (20-25 of each CS presented in pseudo-random order, ‘Post-Learning’) with continued US reinforcement on CS+ trials. During Pre-Learning and Post-Learning trials, contextual cues, consisting of the chamber fabric material and a background odor of either 70% ethanol or 2% acetic acid, were randomly changed across trials.

#### Head-fixed optogenetics

200 *µ*m core, 0.37 numerical aperture (NA) multimode optical fibers were constructed as previously detailed [Kheirbek et al., 2013]. A splitter patch cable (Thorlabs) was used to couple bilaterally implanted optical fibers to a 532 nm laser (50 mW, OptoEngine) for ArchT activation while mice were head-fixed. All cables/connections were shielded to prevent light leak from laser stimuli and matching-color ambient LED illumination was continuously provided in the behavioral apparatus so as to prevent the laser activation from serving as a visual cue. After the 10 second pre-CS period on each trace fear conditioning trial, 10 mW of laser light was continuously delivered through each optical fiber for the entire CS-trace-US sequence.

Experimenters were blinded to subject viral injections. After data collection, mice were processed for histology and recovery of optical fibers. Subjects were excluded from the study if the implant entered the hippocampus, if viral infection was not complete in dorsal CA1, or if there were signs of damage to the optical fiber that could have compromised intracranial light delivery.

#### 2-photon microscopy

For imaging experiments, mice were habituated to the imaging apparatus (e.g. microscope/objective, laser, sounds of resonant scanner and shutters) during the training period. All imaging was conducted using a 2-photon 8 kHz resonant scanner (Bruker). As described in [Danielson et al., 2016], we coupled a piezoelectric crystal to the objective (Nikon 40x NIR water immersion, 0.8 NA, 3.5mm working distance), allowing for rapid displacement of the imaging plane in the z dimension, which permitted simultaneous data collection from CA1 neurons in 2 different optical sections. To align the CA1 pyramidal layer with the horizontal two-photon imaging plane, we adjusted the angle of the mouse’s head using two goniometers (*±*10° range, Edmund Optics). For excitation, we used a 920 nm laser (50-100 mW at objective back aperture, Coherent). Green (GCaMP6f) fluorescence was collected through an emission cube filter set (HQ525/70 m-2p) to a GaAsP photomultiplier tube detector (Hamamatsu, 7422P-40). A custom dual stage preamp (1.4 × 105 dB, Bruker) was used to amplify signals prior to digitization. All experiments were performed at 1.5-2× digital zoom, covering ∼150-200 mm × 150-200 mm in each imaging plane. 2-plane images (512 × 512 pixels each) were separated by 20 *µ*m in the optical axis and acquired at ∼8 Hz given a 30ms settling time of the piezo z-device.

#### Image preprocessing

All imaging data were pre-processed using the SIMA software package [Kaifosh et al., 2014]. Motion correction was performed using a modified 2D hidden Markov model [Dombeck et al., 2007, Kaifosh et al., 2013] in which the model was re-initialized on each plane in order to account for the settling time of the piezo. In cases where motion artifacts were not adequately corrected, the affected data were discarded from further analysis. We used the Suite2p software package [Pachitariu et al., 2017] to identify spatial masks corresponding to neural region of interest (ROIs) and extract associated fluorescence signal within these spatial footprints, correcting for cross-ROI and neuropil contamination. Identified ROIs were curated post-hoc using the Suite2p graphical interface to exclude non-somatic components.

ROI detection with Suite2p is inherently activity-dependent, and so for each session, we detected only a subset of neurons that were physically present in the FOV. Once signals were extracted for all sessions, we registered ROIs across each session as follows. We first chose the session with the largest number of detected neurons as the reference session, and then computed an affine transform between the time-averaged FOV of all other sessions to the reference. Transforms were visually inspected to verify accuracy. Using these transforms, we processed each session serially to register ROIs to a common neural pool across sessions. For a given session (referred to now as the current session), the calculated FOV transform was applied to all ROI masks to map them to the reference session coordinates. We calculated a distance matrix (using Jaccard similarity) that quantified the spatial overlap between all pairs of reference and current session ROIs. We then applied the Hungarian algorithm [Kuhn, 1955] to identify the optimal pairs of reference and current ROIs. All pairs with a Jaccard distance below 0.5 were automatically accepted as the same ROI. For the remaining unpaired current ROIs, pairs were manually curated via an IPython notebook, which allowed the user to select the appropriate reference ROI to pair or enter the current ROI as a new ROI (i.e., not in the reference pool). Any current ROIs whose centroids that were more than 50 pixels away from an unpaired reference ROI were automatically entered as new ROIs, to accelerate the curation. Once all ROIs for the current session were processed (either paired with a reference ROI or labeled as new), the new ROIs were appended to the reference list. The remaining sessions were then processed serially in the same fashion, where the reference ROI list is augmented on each step to include additional ROIs that were not presented in any previously processed session. Once all sessions were processed, this process yielded a complete list of reference ROIs and their identity in each individual session. As a final step, the reference ROIs were warped back to the FOVs of each individual session via affine transform and ROIs that fell outside the boundaries of any session FOV were discarded, so that all analyzed neurons were physically in view for all sessions. Inbound reference ROIs that were not present in individual sessions were assumed to be silent in subsequent analysis.

### Neural data analysis

#### Event detection

All fluorescence traces were deconvolved to detect putative spike events, using the OASIS implementation of the fast non-negative deconvolution algorithm [Friedrich et al., 2017]. Following spike inferencing, we discarded any events whose energy was below 4 median absolute deviations of the raw trace. This avoided including small events within the range of the noise, which could artificially inflate activity rates and correlations between neurons.

Given the dominant sparsity of activity, we then discretized ROI trace to indicate whether an event was present in each frame. Trials for each experiment were collected over the span of several days. Consequently, we found that discretization was necessary to prevent variations in imaging system parameters from exerting undue influences on the analysis, as this could introduce arbitrary variance in the scale of calcium events across sessions.

#### Decoding

All classifiers were support vector machines (SVM) with a linear kernel, using the implementation in scikit-learn [Pedregosa et al., 2011]. For cross-validation, data were randomly divided into non-overlapping training and test sets (75/25% split). This procedure was repeated 100 times for each classifier with random training/test subdivisions and reported as the average across cross-validation folds. Trials were balanced by subsampling the overrepresented class, and all decoding results were compared against a null distribution built by repeating the analyses on appropriately shuffled surrogate data, which controlled for the effects of finite sampling. This is particularly important for fear learning paradigms such as ours, where trial counts are very limited.

#### Decoding elapsed time

We designed a decoder to predict the elapsed time during each trial, in order to assess whether there were consistent temporal dynamics in the neural data during the experiment, such as sequences of time cells. Time decoding was done separately for CS+ and CS-trials, and for Pre- and Post-Learning trial blocks, to assess differences between cues and over learning. We analyzed data during both the tone and trace period, for a total of 35 seconds on each trial. We averaged each neuron’s activity in non-overlapping 2.5 second bins, so that each trial was described by a sequence of 14 population vectors of activity in time.

Our time decoder uses a 1 vs 1 approach through an ensemble of linear classifiers. For each pair of time bins in the experiment, we train a separate classifier to distinguish between population vectors of activity that came from those two time bins. For comparing 14 time bins, this results in a set of 91 binary classifiers trained on all unique time comparisons [Bakhurin et al., 2017, Cueva et al., 2019]. We first evaluated the performance of the individual classifiers by testing their ability to correctly label time bins from held out test trials, where in this analysis each classifier is tested only on time bins from the trial times that it was trained to discriminate. This result is presented as a matrix in Figure 2, and demonstrates the linear separability of any two points in time during the task. We then used the decoder to perform a multi-class time prediction analysis. Here each population vector in a held-out test trial is presented to all classifiers, which “vote” on what time bin the data came from. We take the decoded time to be the time bin with the plurality of votes, and repeat this procedure across all time bins in each test trial to decode the passage of time.

For all time decoding analyses, we compare the classification accuracy or prediction error to a null distribution, which we calculate by repeating these analyses on 1000 surrogate data sets, where for each trial independently, we randomly permute the order of the time bins. This destroys any consistent temporal information across trials, while preserving the average firing rates and correlations between neurons within each trial.

#### Decoding task variables from instantaneous firing rates

We also used a population decoding approach to assess the times in the experiment during which there was significant information about the stimulus identity in the neural data. On each trial, we averaged the activity of each neuron in non-overlapping 1 sec bins. We then trained a separate linear classifier on each time bin to predict whether the population vector came from a CS+ or CS-trial. Classifiers were cross-validated as above on held-out test data. We similarly compared the classification accuracy for trial information to a null distribution, where here we repeated the decoding analysis on 1000 surrogate data sets where the CS trial label was randomly shuffled.

#### Decoding from average firing rates

We similarly assessed our ability to decode task information from the average firing rates of the neurons within a set trial period on each trial. This procedure is identical to the one outlined above, and we used it to assess our ability to decoder the CS identity during the Pre-CS (−5 to 0 sec), CS (0 to 20 sec), Trace (20 to 35 sec), US (35 to 40 sec), and Post-US (40 to 170 sec) periods of the trial (Fig. 4B), as well as during the CS and trace periods combined (0 to 35 sec, 4A).

We additionally assessed how decoders learned at one time period in the task generalize to activity observed in other time periods, by constructing a cross-time period decoding analysis. Here on each cross-validation fold, we trained different decoders to predict CS identity from the activity during each trial period separately. Then on the held-out test data, we tested each decoder not only on the activity from the time period on which it was trained, but on the activity of all other time periods as well (e.g. train on CS, test on trace). The result is a matrix of pairwise trial period comparisons, where the columns indicate the time period used for training the classifier and the row indicate the time period for testing. These comparisons are not necessarily symmetric (e.g., we find that CS-period decoders can be used to predict the cue when tested on trace-period activity better than the reverse).

#### Probabilistic decoding

We additionally assessed our ability to decode the CS identity using a Bayesian approach (see S3). The cross-validation and significance methods were identical to the SVM analysis described previously. For every fold, we used a multinomial naïve Bayes classifier to predict the CS cue from the event counts in a given time bin.

#### Selectivity index analysis

To assess CS-selectivity at the level of single neurons, we computed a selectivity index as:

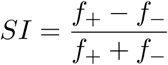

where *f*_+_ and *f*_−_ are the average activity of the neuron from 0-35 sec on CS+ and CS-trials respectively. This yields an index bounded between +1 (all activity during CS+ trials) and −1 (all activity during CS-trial). Similar to the decoding analysis, we compared this selectivity index to those calculated from 1000 surrogate datasets where the trial type labels were randomly shuffled, which controlled for spurious firing rate differences attributable to small numbers of trials. We computed these scores separately for Pre- and Post-Learning trials, and quantified the fraction of cells active during that trial type which showed significant CS-selectivity, determined by calculating a p-value from the observed SI relative to its shuffle distribution. For the regression to population decoder weights shown in Fig. 4E, we z-scored each SI relative to its shuffle distribution’s mean and standard deviation.

#### Ensemble overlap analysis

We measured the similarity in the active set of neurons between a pair of trials by computing the Jaccard similarity index, which for two sets is given by the size of the set intersection divided by the size of the set union (Fig. 5). Intuitively, if the sets of active neurons overlap completely, the set intersection and union are equal and the index is 1, while if trials recruit orthogonal sets of neurons, the intersection and thus the index are 0. This metric is biased by the fraction of active neurons on a given trial; if two trials randomly recruit 50% of neurons, the Jaccard similarity will tend to be higher than if they randomly recruited 10% of neurons, as the former will tend to have more overlapping elements purely by chance. To control for differences in activity rates across trials, we generated 1000 surrogate scores for each trial pair where we recomputed the index between two binary vectors whose elements were randomly assigned as active or inactive to match the fraction of active neurons in the real trials. The observed index was then z-normalized relative to this distribution, to quantify the population similarity beyond that expected from random recruitment of the same number of neurons.

#### Sequence score

For analysis of neural sequences during initial Learning trials, we detected the latency to peak firing rate for each neuron that was active on at least 2 Learning trials. We compared this firing order between all Learning trial pairs via Spearman’s rank correlation, and assigned a sequence score as the average pairwise rank correlation between trials. This analysis was done separately for CS+ and CS-trials. To assess significance, sequence scores were compared to those calculated from 1000 surrogate datasets, where for each neuron, its activity trace was randomly permuted on each trial independently to randomize the temporal ordering between cells.

---

**Figure S1.**
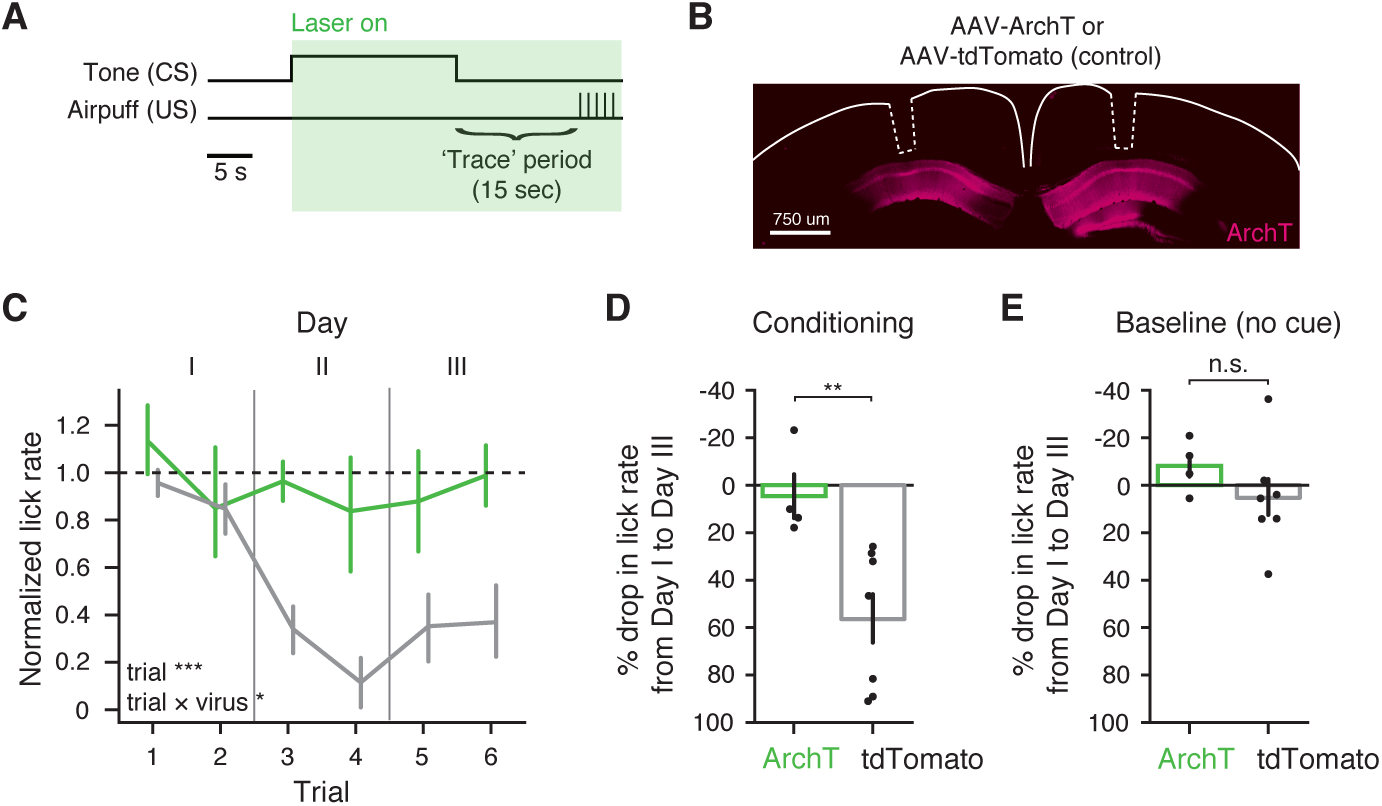
Head-fixed auditory trace fear conditioning is dependent on dorsal CA1 activity. **A:** Schematic of behavior protocol. On each trial, animals are exposed to a 20 second auditory tone (CS) followed by a 15 second trace period. Following the trace period, airpuffs are delivered to the animal’s snout (US). Animals are water deprived and operant rewards are available through each trial. Learned fear is measured via suppression of the licking response. On all conditioning trials, laser stimulation is delivered throughout the tone, trace, and airpuff periods. Animals underwent two conditioning trials per day over the course of 3 days. A baseline trial of the same duration was collected prior to each day’s conditioning trials, where reward was available without presentation of the CS or US. **B:** Left: schematic of optic fiber implants. Mice received bilateral injections in dorsal CA1, with either rAAV1/2-*Syn-ArchT* or rAAV1/2-*Syn-tdTom*, to express inhibitory opsin or control fluorescent protein constructs, respectively. Optic fiber implants were then positioned above the dorsal hippocampus in each hemisphere. Right: *post hoc* confirmation of viral expression and optic fiber placement (dotted lines) in an example mouse. **C:** Normalized licking behavior of *ArchT* and *tdTomato* mice in response to the CS cue for all US-paired trials. Laser stimulation prevents learning only in the *ArchT* group (linear mixed effect model with fixed effects of virus and trial number, and random effect of mouse, significant effects are inset). **D:** Summary of behavioral impact of CA1 inhibition, calculated as the % change in normalized lick rate from day I to III (paired t-test). **E:** As in **D**, but calculated from the baseline (no CS/US presentations) trials on day I and III. Licking behavior is not affected by optogenetic manipulations in the absence of task stimuli. Data are presented as mean *±* s.e.m. \**p <* 0.05, \*\**p <* 0.01, \*\*\**p <* 0.001

**Figure S2.**
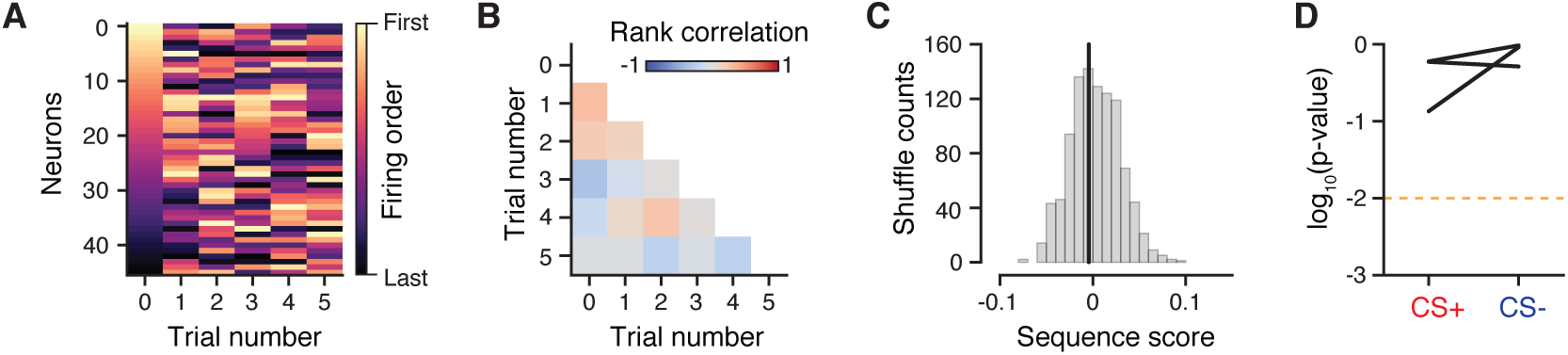
Sequential activity is not reliable during initial learning trials. **A:** Neural sequences during the initial Learning trials, for an example mouse during CS+ trials. Each trial shows the order of neurons’ peak firing time, sorted by firing order on the first trial. Only neurons that were active in at least 2 learning trials are shown. **B:** Rank correlation of peak firing rate times between Learning trials, for the example mouse shown in **A**. **C:** Sequence score (average pairwise trial correlation) for **B**. Grey: null distribution of sequence scores generated by randomly shuffling neuron firing times on each trial. **D:** Summary of learning trial sequence scores for all mice. Neural sequences are not significantly different from random across trials for either trial type.

**Figure S3.**
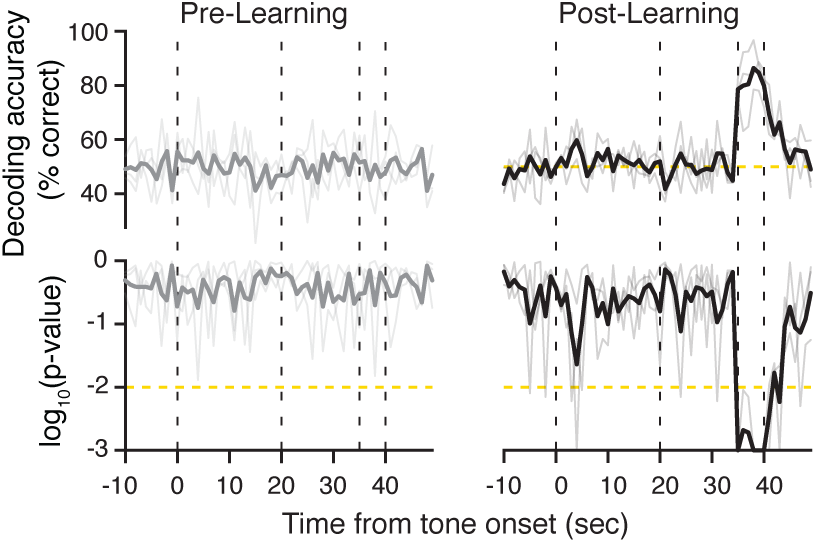
Bayesian decoding of CS identity. We repeated the analysis of decoding accuracy for CS identity shown in Figure 3, here using a naive Bayesian classifier rather than the linear SVM used throughout the paper. The results remained unchanged for both Pre- and Post-Learning trials.

